# Structural development of the human fetal kidney: new stages and cellular dynamics in nephrogenesis

**DOI:** 10.1101/2023.03.24.534074

**Authors:** Anika Schumacher, Tri Q. Nguyen, Roel Broekhuizen, Martijn van Griensven, Vanessa LaPointe

## Abstract

Research on the ultrastructural development of the kidney is limited, and research on rodent kidneys prevails. Yet, large differences between rodent and human nephrogenesis exist and therefore translation between species is not desirable. At the same time, there is an increasing demand for human research, in addition to assessing the potential of novel therapies such as renal organoids. We therefore generated an interactive atlas of large transmission electron microscopy tile scans of first trimester human kidneys, specifically Carnegie stage 20 until post-conceptional week 12. Analysis identified key ultrastructural features of proximal and distal progenitor cells such as cell shape, microvilli and luminal budding in the renal vesicle. Regarding glomerular development, we identified a new W-shaped body stage and three distinct sub-stages of the well-known capillary loop stage. Chromatin organization, nuclear shape and location were used to describe tubule cell identity and maturity, indicating a specific order of tubular maturation. The greatest congruence with adult tissue was seen in proximal tubules and the least in distal tubules. Finally, cytoplasmic glycogen depositions in collecting duct cells, which are absent in adult tissue, were found to be an early feature distinguishing distal tubules from collecting ducts as well as differentiating cortical from medullary collecting ducts. The findings of this research provide new fundamental insights for researchers who aim to understand and recreate kidney development.

## Translational Statement

The findings of this study create more detailed understanding for healthy kidney nephrogenesis. Pathologists and other physicians are currently unaware of the normal ultrastructural features of the human embryonic and fetal kidneys and our findings provide more knowledge in this area. This knowledge will lead to better recognition and understanding of incorrect development and assessment of the effect of specific drugs on nephrogenesis. Aberrant development could then be modelled in kidney organoids for further understanding of the potential cause. Furthermore, our data can aid developmental biologists and tissue engineers in understanding and recreating human kidney development from stem cells and thereby our data can contribute to the improvement of kidney organoids for future transplantation.

## Introduction

Knowledge on the (ultra-)structural development of human embryonic and fetal kidneys is limited and the available microscopy data are largely low in resolution. This dearth of information is partly due to difficulties in obtaining human fetal kidneys to study nephrogenesis. Consequently, researchers who aim to understand human nephrogenesis or need a human reference for lab-grown tissues rely on data from rodent kidneys. However, rodent kidneys are known to develop differently compared to human kidneys.

There are important interspecies differences in ontogeny, as extensively discussed previously.^1-5^ First, kidney development occurs on different timelines. Human kidneys develop within 31 weeks to fully mature and functional kidneys before birth, while mouse and rat kidneys develop in 23 and 18 days, respectively, of which 7 and 8–11 days are after birth.^6^ Second, human kidneys are multi-lobed (4–18 lobes) while rodent kidneys have a single lobe. Third, the number of nephrons^6^ and the glomerular size relative to the total kidney weight differ significantly between species, where for instance mice have a much smaller ratio than humans^1^. Fourth, on the ultrastructural level, differences in brush border lengths of the different proximal tubule segments, the cellular shape of inner medullary collecting duct cells, and the prevalence of peroxisomes in the straight and convoluted proximal tubule have been described.^7^ Last but not least, functional differences are known to exist, for instance in terms of drug clearance.^8^ Additionally, functional kidney development is known to lag behind anatomic maturation in all species^9^, further emphasizing the need for a better understanding of the development of human kidneys. For these reasons, rodent tissue cannot be the ideal reference for human kidneys.

The ultrastructure of human developing kidneys, as visible by electron microscopy, has been analyzed to limited extents. Developing human glomeruli have been studied most extensively, compared to other nephron segments. Early research determined the location and morphology of microvilli and cilia and the overall structure of the glomerulus, revealing that the ultrastructural features are preserved when the explant is kept in culture.^10^ Other studies quantified metrics such as the volume density of mitochondria and lysosomes, and the surface density of the basolateral membrane^11^, as well as the location of intracellular contractile filaments in podocytes and mesangial cells^12^. Later on, given the variety of diseases involving glomerular integrity, tight junctions in foot processes gained interest and slit filtrate proteins were determined.^13^ The most recent and comprehensive study on the glomerular structure of 10- to 22-week-old human kidneys visualized the 3D shape of developing podocytes by scanning electron microscopy (SEM) and correlated this data with ultrastructural characteristics.^14^ In contrast, the study of human tubular development has been largely limited to SEM studies of gross morphology^15^ and ultrastructural information is lacking. Clearly, more ultrastructural features can be identified with today’s improved microscopy and the increased need for understanding human nephrogenesis makes this an important topic to address.

Our aim was to generate an atlas of high-resolution data obtained with transmission electron microscopy (TEM) of the various stages of human nephrogenesis of the entire kidney. We selected age groups Carnegie stage 20 (CS20) until post-conceptual week (PCW) 12, the latter comparable to stem cell–derived constructs such as kidney organoids^16^. Our novel ultrastructural data provide important insight into human nephrogenesis that can aid developmental biologists and tissue engineers in recreating complete and functional kidney tissue.

## Methods

### Patient tissue

The human embryonic and fetal material was provided by the Joint MRC/Wellcome Trust (grant# MR/R006237/1) Human Developmental Biology Resource (www.hdbr.org). The medical files were screened to exclude kidney diseases. After excision, the kidneys of each embryo/fetus were longitudinally cut in half, if possible, of which one half was fixed for histology and the other half was fixed for TEM. The kidneys (summarized in Table 1) were confirmed normal by a pathologist (T.N.) and technician (R.B.) of the University Medical Center Utrecht (the Netherlands) and were subsequently analyzed using TEM. A minimum of two cubes were viewed per patient. However, the CS20 and week 8 kidneys were so small that either the whole or nearly half kidney was seen in two blocks.

**Table 1:**
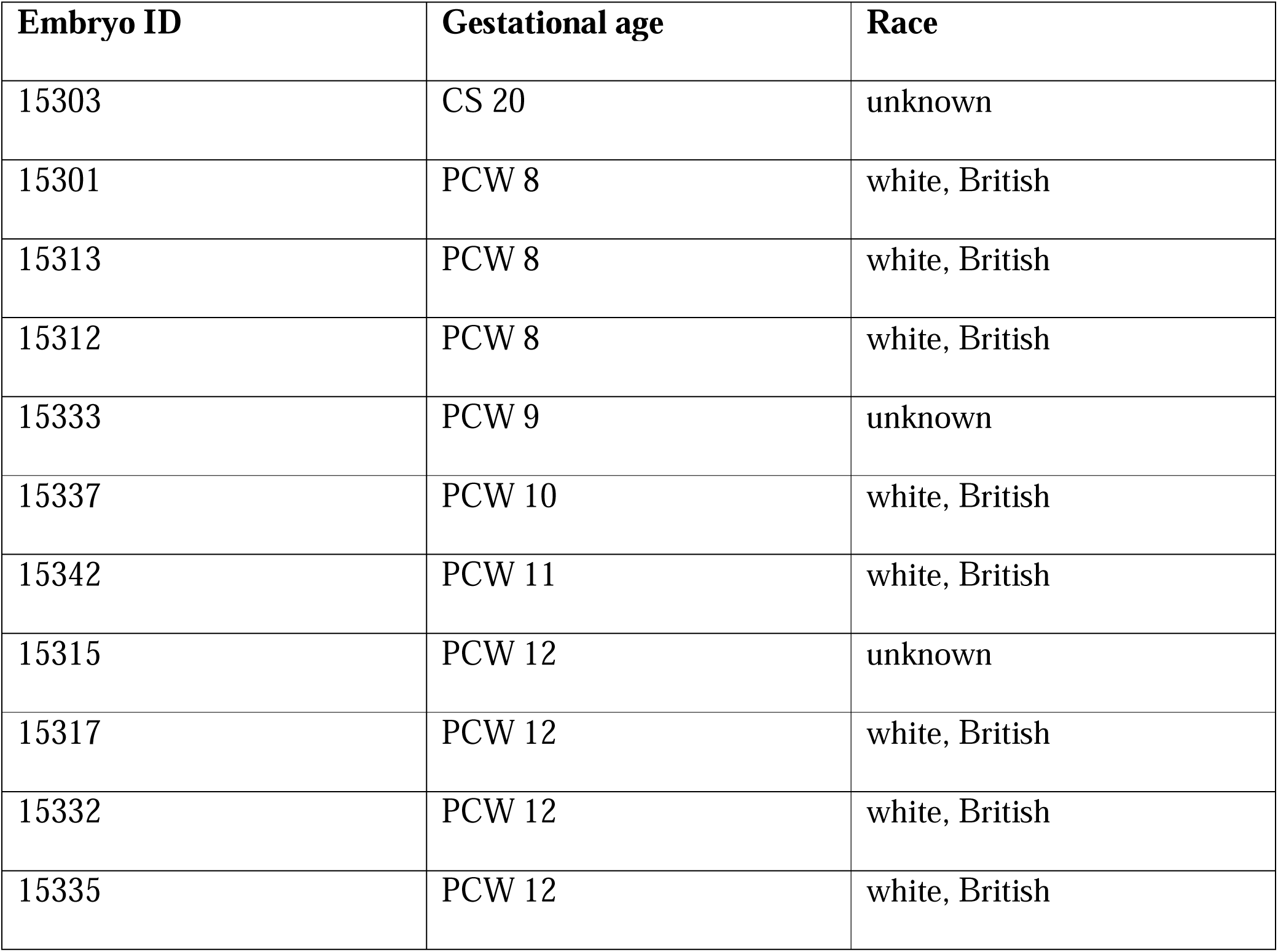
Patient information. Carnegie stage (CS), post-conceptional week (PCW).

### Histological staining

The kidneys were immediately fixed after explantation with 4% formalin overnight at 4 °C and embedded in paraffin for histological staining. The tissue was cut into 3 µm slices. Periodic acid-Schiff (PAS) and hematoxylin and eosin (H&E) staining were performed using standard procedures (Dako, Glostrup, Denmark) for the assessment of normal structures.

### Tissue embedding and processing for TEM

Tissue embedding and processing was performed according to standard procedures described in detail in the Supplementary Methods. Briefly, the kidneys were cut into 1 mm^3^ cubes while immersed in Karnovsky’s fixative. After washing with 0.1 M cacodylate buffer, the samples were postfixed with 1% osmium tetroxide and 1.5% potassium ferricyanide. After washing and dehydration, the samples were embedded in Epon. The region of interest (ROI) was determined in 1 μm sections stained with toluidine blue (Figure S1). Ultrathin sections of 60nm were then cut on a Leica UC7 ultramicrotome and stained with 2% uranyl acetate and Reynold’s lead citrate.

### TEM imaging

Next, the sections were observed on a Tecnai T12 electron microscope equipped with an Eagle 4k × 4k CCD camera (Thermo Fisher Scientific). To target the ROI, full-section scans were made using the EPU software (1.6.0.1340REL, FEI). Subsequently, the ROI was imaged in tiles at 2900× magnification using the EM Mesh software (version 6). Data were stitched, uploaded and annotated using Omero and PathViewer. The data are available as open source on the Image Data Resource (https://idr.openmicroscopy.org) under accession number idr0148.

### FIB-SEM acquisition and 3D rendering

The ROI was targeted by cutting the block with a Leica UC7 ultramicrotome. The blocks were mounted and then coated with a thin layer of heavy metal to prevent charging. FIB-milling was done at a 0.5 nA beam current, where gallium ions were accelerated to 30 kV. Serial block-face images were taken every 10 nm with a backscattered electron detector at an acceleration voltage of 2 kV and 0.4 nA current. The pixel width was 10 nm and the pixel dwell time was 300 ns. Nuclei were segmented and chromatin thresholded as described in the Supplementary Methods.

## Results

### Early stage nephrogenesis is characterized by uteric bud (UB)-guided maturation

The cap mesenchyme (CM) adhered to the ureteric bud (UB) of PCW 8 kidneys in the nephrogenic zone to induce differentiation of mesenchymal progenitors into pretubular aggregates (PTA) and later into renal vesicles (RV) (Figure 1a). The well-known dynamic nature of the CM^19^ could be seen in the irregular cell shape and loose intercellular connections to the ECM (Figure 1b,c). In contrast, the UB consisted of tight epithelial layers (Figure 1d) with apoptotic cells located in the lumen of the UB tip (Figure 1e). Large glycogen depositions were located both apically and basally throughout the UB cell layers (Figure 1f, Figure S2). The more differentiated PTA cells showed the most epithelial-like morphology as characterized by polarized, tightly aligned cells with basally located nuclei and were loosely attached at the medial side to the UB (Figure 1g,h). In contrast, the PTA cells located laterally, namely furthest from the UB, still possessed mesenchymal-like irregular shapes with loose cell–cell contact and no polarity (Figure 1i).

**Figure 1:**
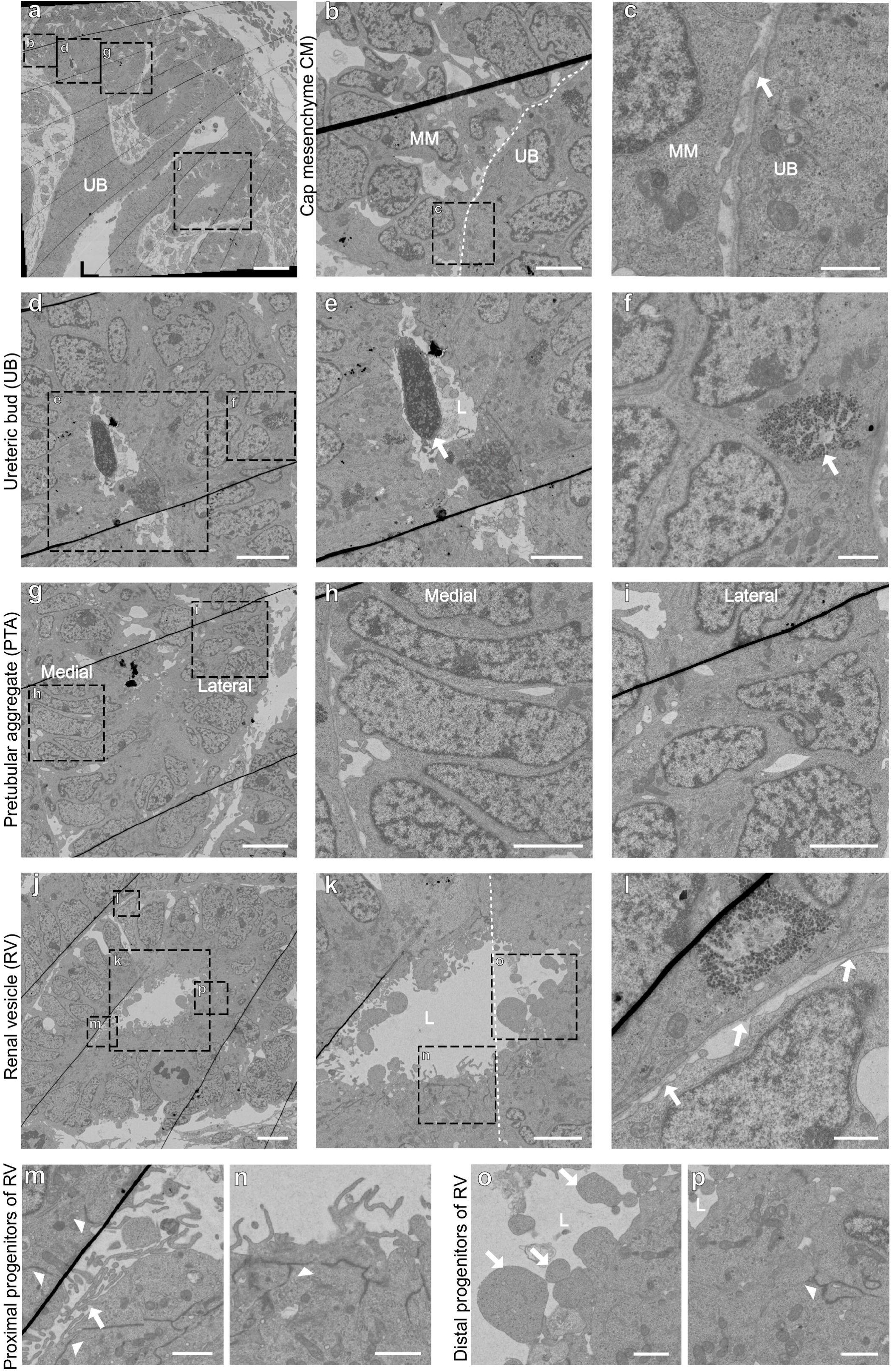
Nephrogenesis in PCW 8 tissue (patient ID 15301) indicates marginal cellular maturation. (a) An overview of early nephrogenic stages: branched ureteric bud (UB) in contact with the cap mesenchyme (CM), forming a pretubular aggregate (PTA) and renal vesicle (RV). (b) Irregularly shaped CM and organized epithelial UB confirm the known dynamic interaction. The dotted white line indicates the border between CM and UB where (c) the CM adhered to the UB by extracellular matrix (white arrow). (d) The UB tip features (e) apoptotic luminal cells (white arrow) potentially due to lumen formation and (f) large basally located glycogen (Gly) depositions (white arrow). (g–i) Cells of the PTA have distinct shapes indicating regional maturation. (h) Epithelial-like cell shape is prominent closest (medial) to the UB, while (i) a mesenchymal-like, irregular shape was seen in lateral cells. (j) Cells of the RV have distinct apical patterning: (k) Proximal progenitors are located left, and distal progenitors right, of the white dotted line. (l) Proximal progenitors adhering to the basement membrane of the UB (white arrows). (m–n) Proximal progenitors possess microvilli (white arrow), fewer buds, and apical tight junctions (white arrowheads). (o) Apical budding (white arrow) of the distal progenitors is potentially a mechanism of luminal mitosis. (p) Large tight junctions (white arrowhead) are distanced from the lumen (L). Scale bars: a: 20 µm; d,g,j: 5 µm; b,e,h,i,k: 3 µm; c,f,l-p: 1 µm.

At the next developmental stage, the RV, a lumen had formed (Figure 1j). Similar to the PTA, the RV was loosely connected via ECM proteins to the UB and the most epithelial-like cell shape could be found medially next to the UB (Figure 1j,l). The lumen showed two interesting features. First, in the region known to contain the proximal progenitors (Figure 1k; left of the dotted line), cells possessed microvilli (white arrow) and tight junctions were clearly located apically (Figure 1m,n; white arrowheads). Second, throughout RV, the RV cells were budding into the lumen (Figure 1o; white arrows), and distal progenitors (Figure 1k; right of the dotted line) had tight junctions that were not adjacent to the lumen (Figure 1p; white arrowhead). Interestingly, microvilli on distal progenitors were rare.

### Novel stages of glomerular development

At the s-shaped body (SSB) stage, in which the intra-glomerular capillaries had yet to develop and there were only endothelial cells invading the cleft (Figure 2a), ultrastructural analysis of human fetal and embryonic kidneys revealed a tightly packed podocyte population connected by tight junctions (white arrowheads) with moderately elongated shapes (Figure 2b). At the tip of the SSB, a small lumen was observed between the podocytes and parietal epithelial cells (PECs), into which small protrusions extended from the podocytes (white arrow) near their apical tight junctions (Figure 2c; white arrowhead). We identified a new stage following the SSB stage, which we term the W-shaped body (WSB) stage for its W-like shape. Characteristics of this stage are the large invasion of endothelium, capillaries and mesangial cells (Figure 2d, white arrow) in consort with a lack of fully formed glomerular capillaries and glomerular basement membrane (GBM). Regional differences in maturity were characterized by the amount of deposited GBM as well as podocyte morphology. When matrix was deposited (white arrow), the endothelium (END) and podocytes (P) were connected, leading to podocyte polarization (Figure 2e–f). In comparison, when little matrix was deposited, the endothelium and podocytes were separated, and the podocytes were not polarized (Figure 2g–h).

**Figure 2:**
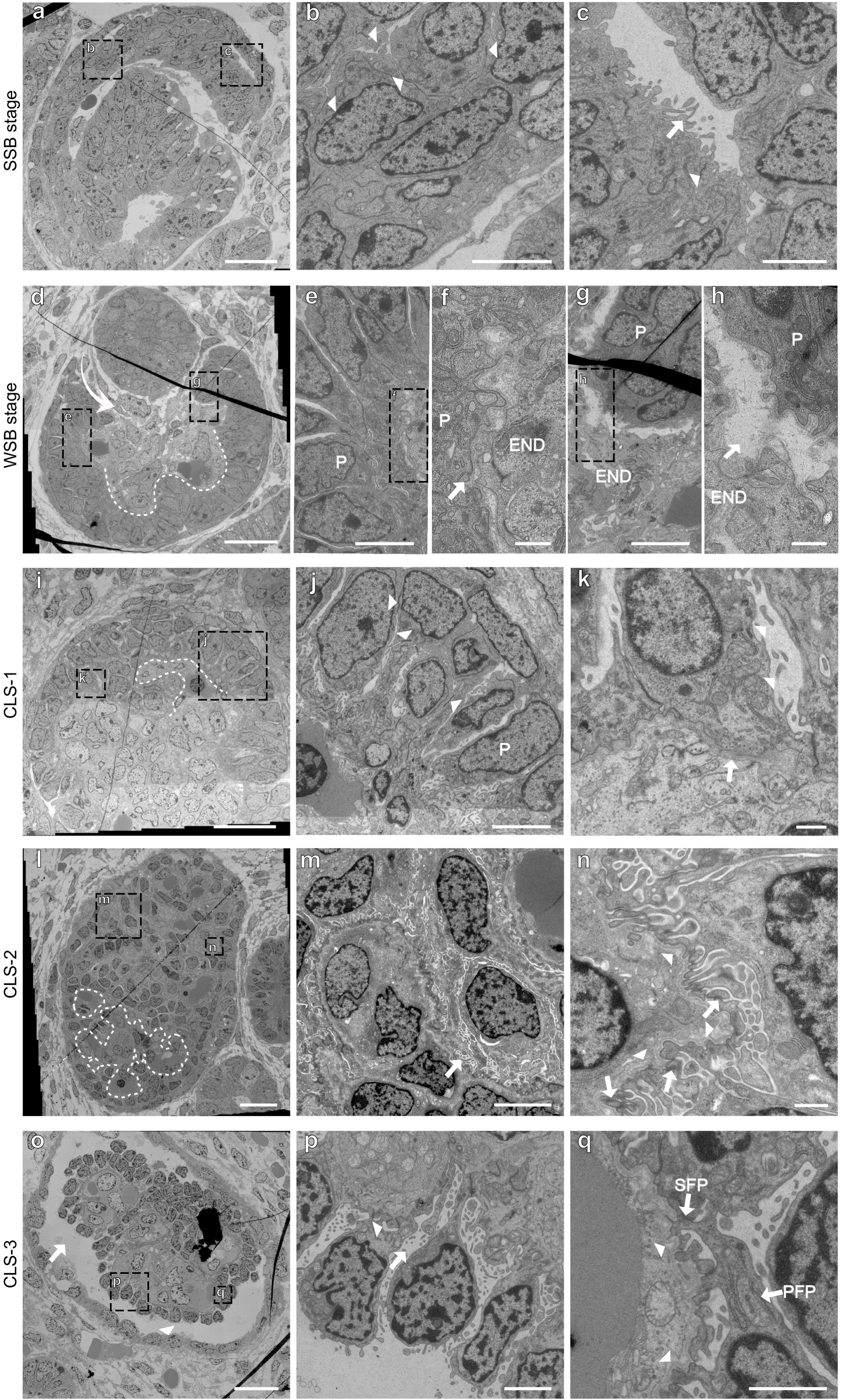
Ultrastructural analysis reveals novel stages of early glomerulogenesis. (a) S-shaped body (SSB) glomerulus (patient ID 15335) shows (b) podocytes of irregular cell shape, which are densely packed and connected by tight junctions (white arrowheads). (c) Protrusions (white arrow) extending from podocytes to parietal epithelial cells (PECs) on the outer SSB tip. Apical tight junctions (white arrowhead) indicate that cells are of epithelial phenotype. (d) A newly identified W-shaped body (WSB)-stage (patient ID 15315) is characterized by major endothelial, mesangial and vessel infiltration (curved white arrow), where regional podocyte maturation is related to the extent of glomerular basement membrane deposited between endothelial cells and podocytes: (e) polarized podocytes (P), where (f) the basement membrane is deposited towards the endothelial cells (END, white arrow), versus (g) unpolarized podocytes where (h) the basement membrane is lacking (white arrow). (i) Capillary loop stage (CLS)-1 features immature capillary loops (white dotted line) (patient ID 15335), (j) polarized podocytes connected at the cell bodies via tight junctions (white arrowheads), (k) an immature GBM (white arrow) and the podocyte microvilli (white arrowheads). (l) CLS-2 has fully formed capillary loops (white dotted lines) (patient ID 15301), (m) polarized podocytes with microvilli (white arrow), (n) foot processes (white arrows) and a complex multilayered GBM (white arrowheads). (o) CLS-3 is the most mature glomerulus of the first trimester, featuring a large Bowman’s space (white arrow), a single layer of parietal epithelial cells (white arrowhead) (patient ID 15342), (p) urinary space in between podocytes (white arrow) and around foot processes (white arrowhead). (q) Foot processes matured into primary foot processes (PFP) and secondary foot processes (SFP) integrated into the glomerular basement membrane (white arrowheads) with urinary space in between. Scale bars: d,i,l,o: 20 µm, b,e,g,j,m: 5 µm, a,c,p: 3 µm, q: 2 µm, f,h,k,n: 1 µm.

To date, the capillary loop stage (CLS) has been described as a single stage of glomerular development in which the glomerular capillaries are formed. However, in-depth morphologic analysis showed that developmentally distinct CLSs could be distinguished that we termed CLS-1, CLS-2, and CLS-3. CLS-1 featured a continuous GBM with endothelium and podocytes attached to either side (Figure 2i). The podocytes were polarized in this stage, with basal nuclei and tight junctions between cell bodies (Figure 2j; white arrowheads) and primary foot processes connected to the GBM (Figure 2k; white arrow) and interconnected through tight junctions. The presence of microvilli on podocytes was observed for the first time (Figure 2k; white arrowheads). The next stage, CLS-2, featured fully formed glomerular capillaries, which were rich in blood cells (Figure 2l, white dotted lines). Podocytes adhered to the multilayered GBM (Figure 2m, white arrowheads) that possessed secondary foot processes with tight junctions (Figure 2n, white arrows), forming the filtration barrier. The final stage, CLS-3 (Figure 2o), was characterized by a large Bowman’s space (white arrow) and a single layer of PECs (white arrowhead). Mature characteristics were present (Figure 2p), such as large urinary spaces and microvilli between podocytes (white arrow) and urinary space between foot processes (white arrowhead). The GBM was curled, with integrated secondary foot processes (SFP) extending from primary foot processes (PFP) (Figure 2q). The GBM (Figure 2q; white arrowheads) consisted of the lamina densa (dark gray), and on either side, lamina rara externa and lamina rara interna (light gray). All features are summarized in Figure 3.

**Figure 3:**
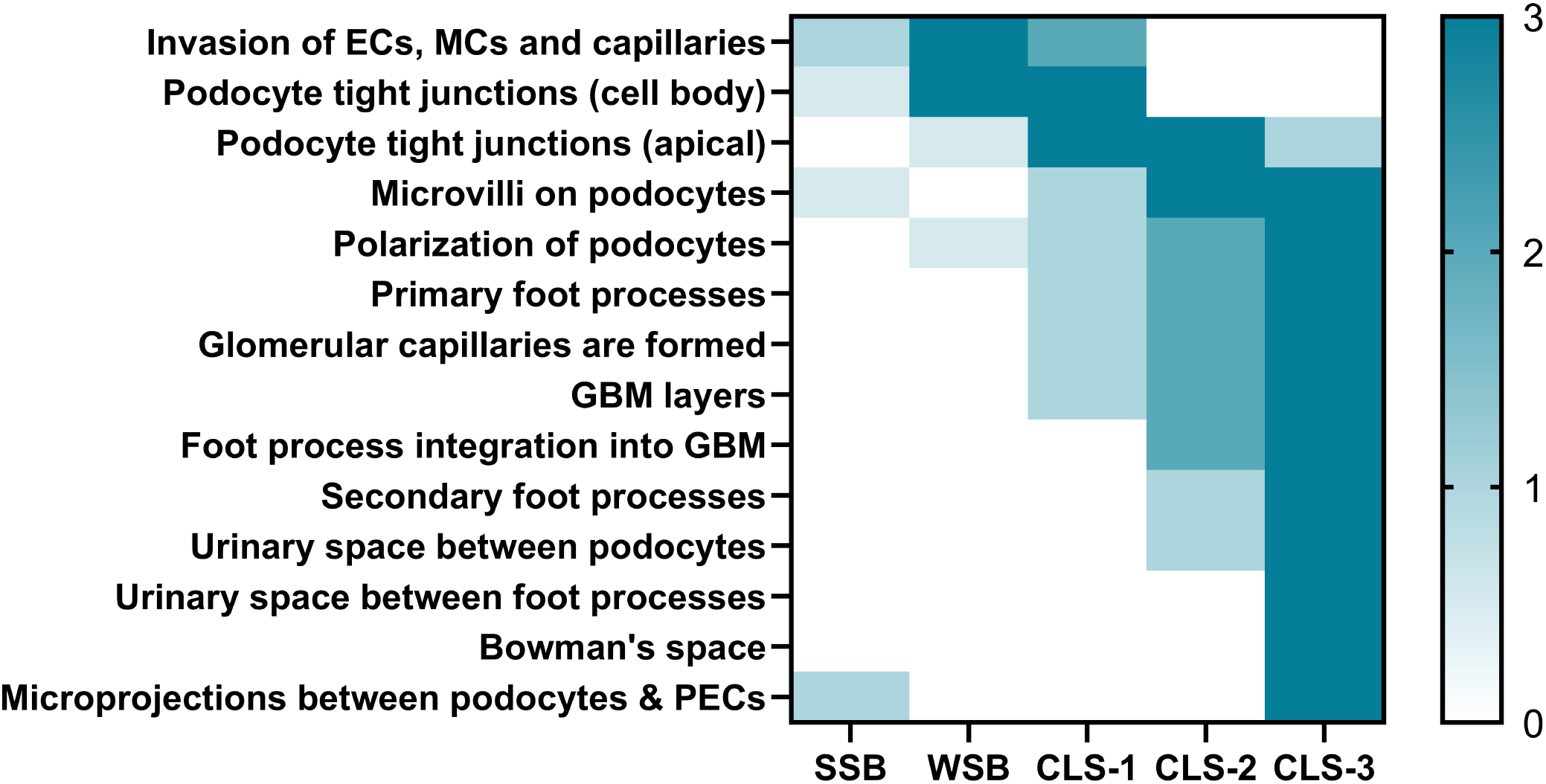
Summary of features of glomerular development from SSB to the most mature CLS-3 in the first trimester. ECs = endothelial cells, MCs = mesangial cells, GBM = glomerular basement membrane, PECs = parietal epithelial cells, SSB = S-shaped body, WSB = W-shaped body, CLS = capillary loop stage. Heat map indicates relative abundance with 0 = not present and 3 = maximum presence.

### Changes of nuclear morphology indicate proximal tubule maturation

To date, knowledge on the structural features of first trimester tubules has been insufficient to distinguish proximal from distal tubules. By comparing the ultrastructure at different ages, we could identify how the features describing mature PTs, such as brush borders, basal round-shaped nuclei, and lysosomes, emerged from immature PTs. Although the increase in maturity between stages corresponds to an increase in PCW, we named these stages 1–3. This is because the kidney matures in a hierarchical manner and therefore the most mature (stage 3) tubule can also be seen in week 9 kidneys towards the medulla. Accordingly, the least mature tubule can be seen adjacent to the nephrogenic zone in week 11 (data not shown).

Stage 1 PTs showed an incomplete epithelial morphology (Figure 4a), since cell–cell contact was predominantly on the apical side, while basally, a mesenchymal-like morphology was still visible. Nuclei were located basally, but were irregular in shape (Figure 4b). Small clusters of microvilli indicated the formation of a brush border (Figure 4c, white arrowheads). In stage 2 PTs, epithelial cell–cell contact was largely established, and the nuclear shape changed to smooth oval–round shapes (Figure 4d,e). The brush border was more developed (Figure 4f) than in stage 1. At stage 3, the PTs had matured further and showed signs of functionality (Figure 4g) in that complete cell–cell contact was established to allow the characteristic active transport (Figure 4h). The nuclei were homogenously round, and electron-dense lysosomes could be detected, particularly in regions close to peritubular capillaries (Figure 4h), indicating increased functionality. The brush border could be seen throughout the whole tubule cross-section and the vacuolar apparatus increased in complexity (Figure 4i). The different tubule stages were compared to previously published rat proximal tubule cross sections, and stage 3 proximal tubules were found to have most congruency. Images of rat tissues were chosen as a comparison since, to our knowledge, no TEM images of adult human proximal tubule cross sections are available in equally high resolution and large field of view.

**Figure 4:**
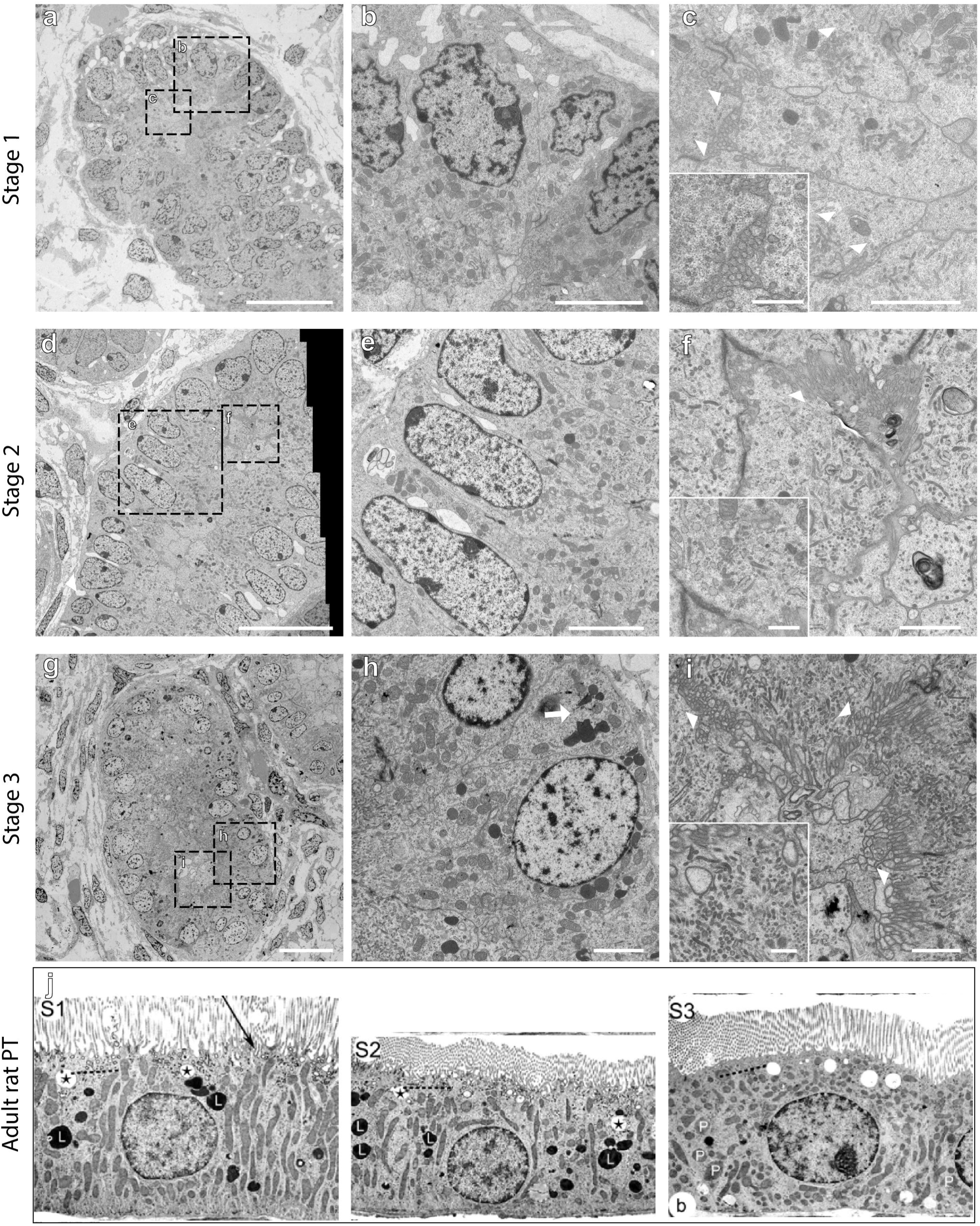
Features of maturing proximal tubules. (a) Immature proximal tubule (patient ID 15333, PCW 9) with (b) basally located, irregularly-shaped nuclei, and (c) an immature apical brush border (white arrowheads). (d) Further matured proximal tubule (patient ID 15337, PCW 10) features smooth, oval nuclei, apically located in (e) PT epithelial cells characterized by increased polarization and cell–cell contact, and (f) increased number of apical microvilli (white arrowhead). (g) The most mature PT stage of the first trimester (patient ID 15342, PCW 11) was characterized by (h) round, basal and light-colored (low electron density) nuclei with dispersed chromatin and the appearance of lysosomes (white arrow), (i) a larger brush border (white arrowheads) with increased vacuolar apparatus (inset) compared to previous stages (insets in c and f). (j) Cross-sections of adult rat proximal tubules with round nuclei and dispersed chromatin. S1-S3 correspond to the different PT segments (adapted from^46^). Scale bars: g: 30 µm, a,d: 20 µm, b,e: 5 µm, c,h: 3 µm, f,i: 2 µm, insets: 1 µm.

Three-dimensional reconstruction of nuclei of both stage 1 and stage 3 proximal tubules confirmed the nuclear morphology seen in TEM (Figure 5). The stage 1 nucleus was highly irregular in shape (Figure 5a,b), while the stage 3 nucleus was homogenously round shaped (Figure 5c,d). Segmentation of electron dense chromatin showed a trend for chromatin reorganization during maturation. A proof of concept quantification of the radial distance of electron-dense chromatin from the nuclear envelope indicated a larger nuclear envelope– bound fraction in the stage 1 nucleus compared to the more mature stage 3 nucleus, while the overall fraction of electron-dense chromatin remained the same (Figure S3).

**Figure 5:**
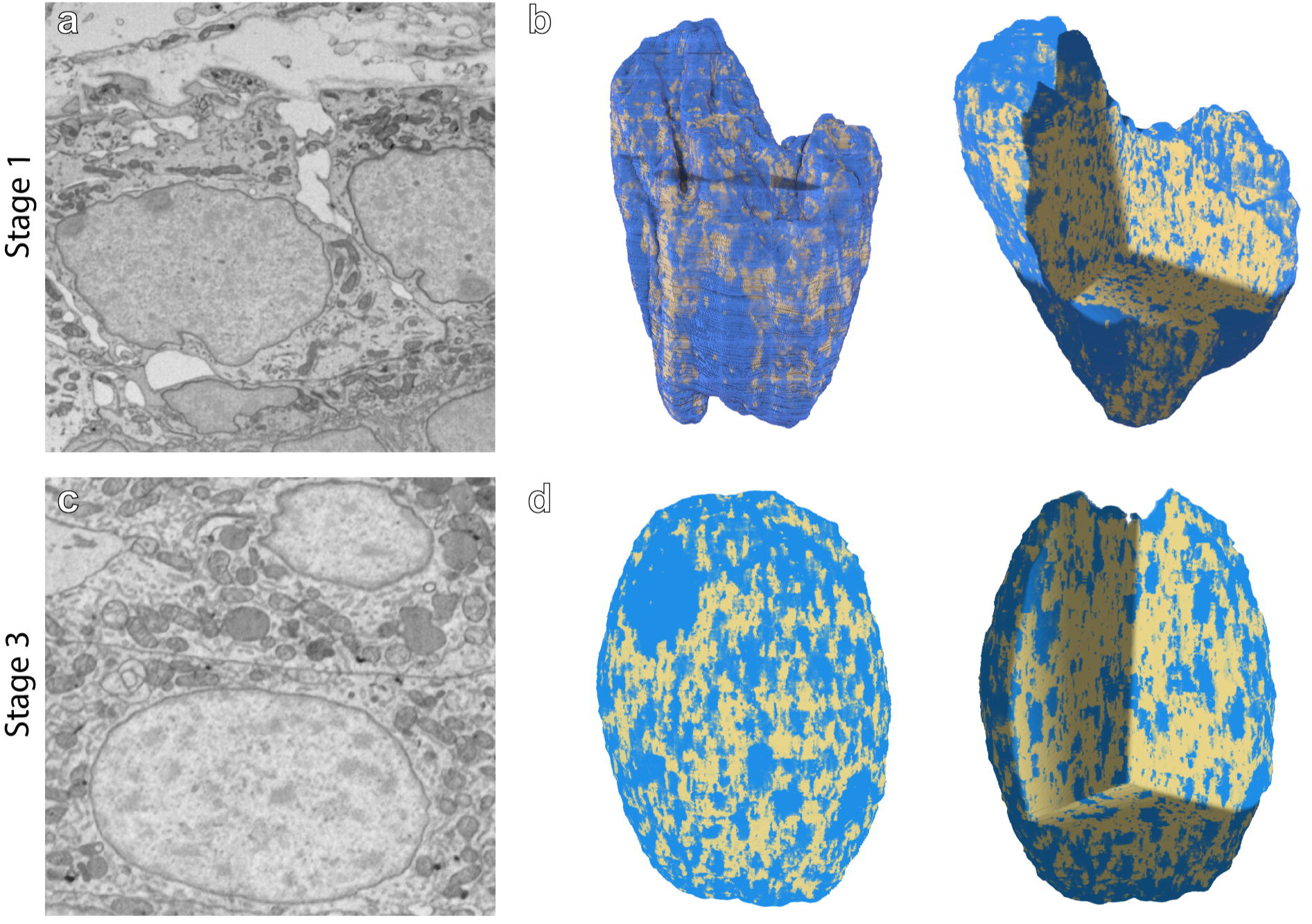
Nuclear reconstruction of developing proximal tubules highlights changes in nuclear shape and chromatin organization with increasing maturity. (a) FIB-SEM slice through middle of a stage 1 (least mature) nucleus chosen for segmentation. (b) 3D rendered stage 1 nucleus (blue) with segmented chromatin (beige) (patient ID 15333). (c) FIB-SEM slice through the middle of a stage 3 (most mature) nucleus chosen for segmentation. (d) 3D rendered stage 3 nucleus (blue) with segmented chromatin (beige) (patient ID 15342).

### Ultrastructural features indicate an order of maturation of tubules

The identification of the most mature tubules based on their location and well-known features of maturity led us to identify less mature tubules and describe the order in which the different tubules matured. As previously explained, these findings are not presented in terms of patient age, however, ages are stated in the caption of Figure 6.

**Figure 6:**
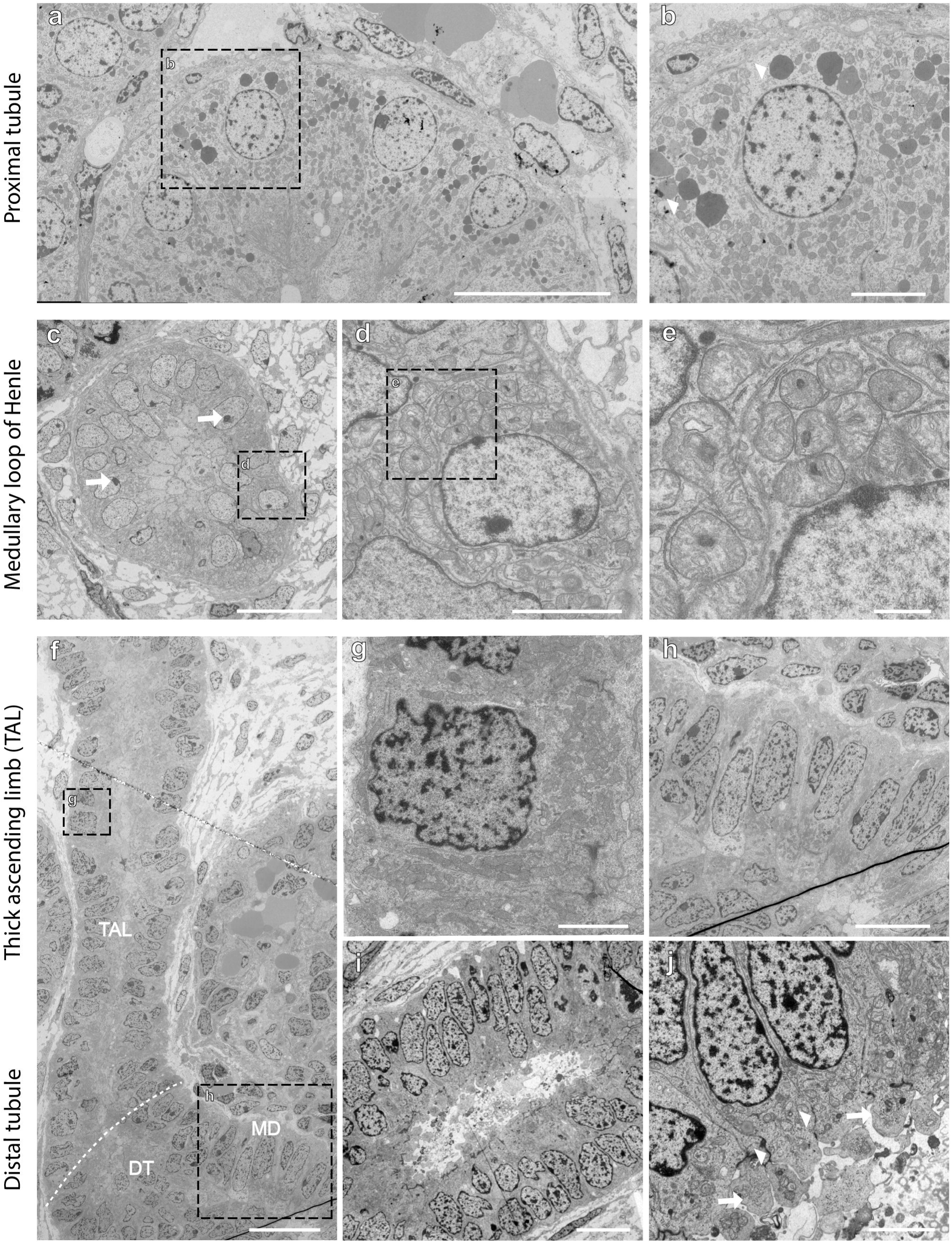
Nuclear shape, electron density, nuclear location and chromatin organization are features to distinguish the most mature developing tubules of the first trimester. (a-b) The proximal tubule features evenly aligned cells with round, light colored (low electron density) nuclei, speckled chromatin, and electron dense lysosomes (white arrowhead) (patient ID 15342, PCW 11). (c-d) The medullary loop of Henle has cells with evenly distributed chromatin and a clearly visible nucleolus (white arrows) and (e) the largest number of mitochondria with well-defined cristae (patient ID 15333, PCW 12). (f) The cortical, thick ascending loop (TAL) of Henle is directly connected to the distal tubule (DCT) after the macula densa (white dotted line) (patient ID 15333, PCW 12). (g) TAL cells are characterized by irregularly shaped nuclei that are basally located with irregular and dense chromatin organization and a large number of mitochondria. (h) The distal tubule possesses aligned macula densa cells (MD), which are comparable to other DCT cells (i-j) with largely oval, centrally located nuclei and irregular chromatin organization (patient ID 15317, PCW 12). Scale bars: a,c,f: 20 µm, h,i: 10 µm, b,d,j: 5 µm, g: 3 µm, e: 1 µm.

Proximal tubules had the most resemblance to adult kidneys (Figure 4, Figure 6a,b), as seen in their homogenous, round nuclear shape and basal location, and the appearance of lysosomes and a brush border. The chromatin was equally dispersed throughout the nucleus, however it was morphologically distinct from adult PTs (Figure 4g–i, Figure 6a,b). The medullary loop of Henle had irregularly shaped nuclei, located both apically and centrally, with a prominent nucleolus (white arrows) and dense clusters of chromatin associated with the nuclear envelope. The nucleus was small compared to the cytoplasm (Figure 6c). The large number of mitochondria (Figure 6d,e) corresponds to the need to sustain oxidative phosphorylation in the less vascularized medulla.^20^ The cytoplasm to nucleus ratio of the cortical thick ascending limp (TAL) was comparable to the medullary LoH with prominent cytoplasm on the apical side (Figure 6f–g). The TAL was similarly rich in mitochondria as the medullary LoH (Figure 6g). The nuclear shape and chromatin organization, however, were distinct from the medullary LoH. Namely, the nuclei were irregularly shaped with dense clusters of chromatin, both dispersed throughout the nucleus and attached to the nuclear envelope, potentially indicating a less mature state than the medullary LoH (Figure 6f,g). Immediately after the TAL, the distal tubule is connected^21^ and characterized by the macula densa with tightly aligned cells with large, elongated nuclei^22^ (Figure 6f,h). The distal tubule had similar characteristics (densely aligned, partially elongated cells with elongated nuclei), but also showed large apical tight junctions and apical budding. The nuclei were irregularly shaped and showed dense clusters of chromatin dispersed throughout (Figure 6i,j). Morphological distinction of proximal tubule, distal tubule and collecting duct were confirmed using specific immunostaining combined with histological PAS staining (Figure S4).

### Features to distinguish cortical and medullary collecting ducts

The cortical collecting duct developed and matured before the mesenchyme-derived nephrons and was therefore characterized by several layers of organized epithelial cells containing large glycogen deposits (Figure 7a; white arrows). Like the UB tip, the collecting duct possessed large cytoplasmic glycogen deposits both apically and basally. Within the UB tip and further towards the medulla, apoptotic cells could be found in the lumen (Figure 7b; white arrow). The UB tip and residual part of the CD tubule had short apical microvilli (Figure 7c; white arrows). Further distally in the CD in direction of the medulla, cells at the apical side were budding into the lumen (Figure 7d; white arrows), connected by tight junctions (Figure 7e; white arrowheads). Medullary CDs lacked this apical budding, but short microvilli persisted (Figure 7g; white arrows). Cytoplasmic glycogen deposits increased in size and number in the medullary CD compared to the cortical CD and were largely located in the basal cell layer (Figure 7f–h; white arrowheads). These findings were consistent from week 8 to week 12, making glycogen deposits a reliable feature to identify first trimester UBs.

**Figure 7:**
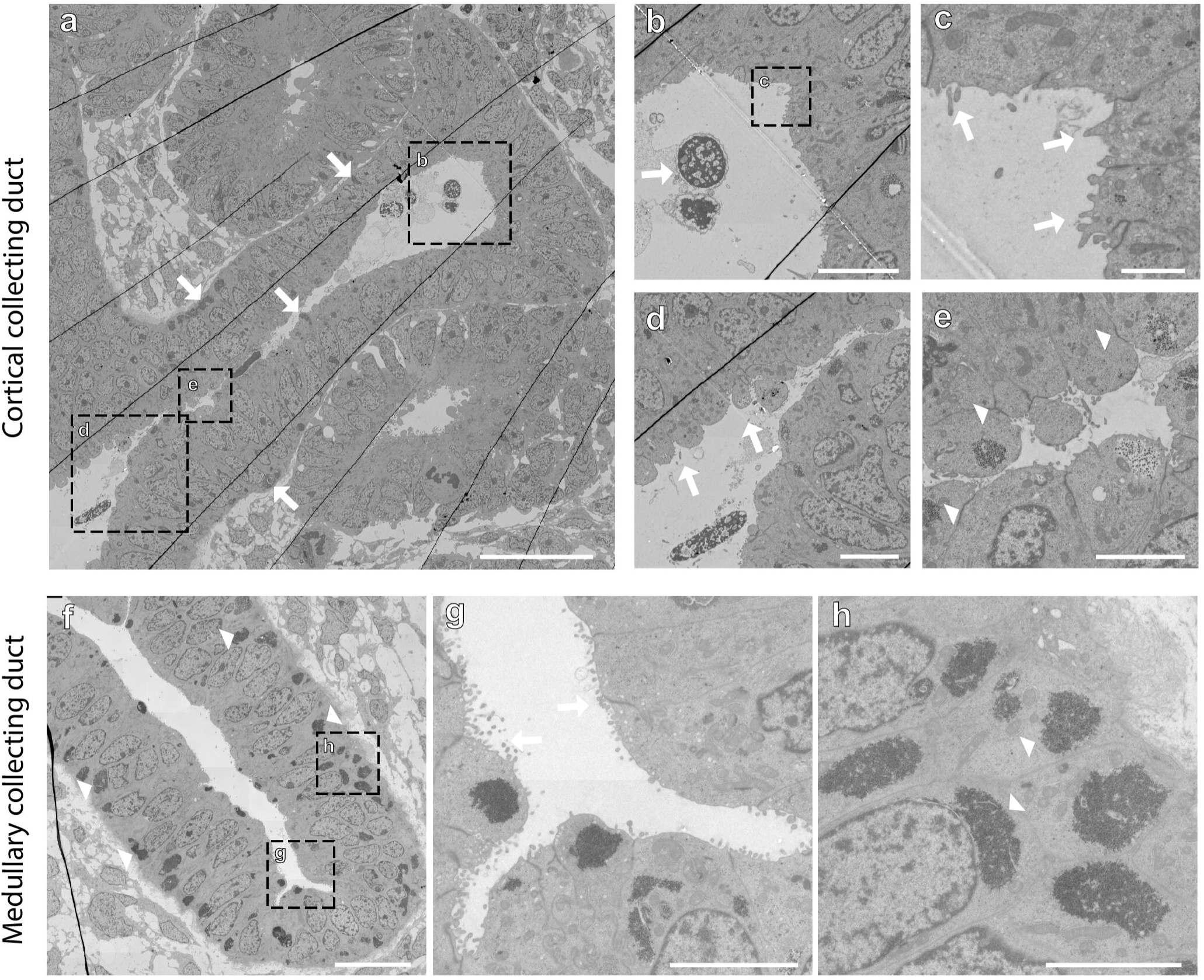
Features of developing medullary and cortical collecting duct (patient ID 15301). (a) Cortical collecting duct (CD) spanning throughout the cortex up to the nephrogenic zone contains large glycogen deposits (white arrows). (b) Apoptotic cells are located throughout the lumen (white arrow). (c) CD cells have short, apical microvilli (white arrows). (d) CD cells further from the renal vesicles, towards the medulla, show apical budding (white arrows). (e) Tight junctions are located between the buds (white arrowheads). (f) Medullary CD is richer in glycogen deposits than the cortical CD (white arrowheads). (g) Short microvilli are located at the apical side (white arrows) and buds are lacking. (h) Excessive glycogen deposits (white arrowheads) are located mostly on the basal side. Scale bars: a,f: 20 µm, b,d,g,h: 5 µm, e: 3 µm, c: 1 µm.

## Discussion

Our aim was to generate a TEM-based atlas of first trimester human nephrogenesis in the metanephric kidney, in which we define features that allow the identification of cell types in immature states and enable their correlation to known functionalities. Our analysis revealed novel features throughout the entire nephron, from nephron progenitors to glomerular and tubular differentiation.

The mammalian kidney is known for its unique and complex morphology, which underlies its functionality. The first blood filtration takes place around week 9 in human kidneys^23^ implying a functional ultrastructure must form in advance of this time. The earliest stage of our study, CS20, was characterized by the close proximity of the CM and UB (Figure S5), indicating the start of nephrogenesis. One week later, at week 8, PTAs and RVs had formed, and the first generation of immature proximal, distal tubule and capillary stage glomeruli developed (Figure S5, Figure 1). The first glomerular generation was comparable to a CLS-3 stage glomerulus and the proximal tubule (PT) comparable to stage 2, as newly defined in this study (Figures 2 and 4). The formation of a CLS-3 stage glomerulus confirmed the early formation of functional ultrastructure and indicated the possible start of glomerular filtration. Nevertheless, it cannot be neglected that a larger sample size per age could potentially show different timings in the emergence of ultrastructural features due to differences in maturation between individuals.

At week 8, features of early cell type specification were determined to distinguish tubule progenitors in the RV stage, close to the nephrogenic zone. The first cell type specification towards proximal and distal progenitors is thought to occur at the RV stage, which has been shown transcriptionally and confirmed by immunofluorescence.^24, 25^ While these populations are histologically indistinct,^26^ our data suggest that ultrastructural differences exist. In contrast to distal progenitors, proximal progenitors possessed short microvilli and apical tight junctions immediately adjacent to the lumen (Figure 1k), as shown in early research on mouse RV.^27^ The presence of microvilli in RVs has not been investigated and therefore their function is unknown. Mechanosensors exist in the brush border of the proximal tubule,^28–30^ however given the absence of a continuous lumen towards the CD until the SSB stage,^24^ and consequently the absence of fluid flow, mechanosensation to sense flow in the RV is unlikely. Nonetheless, luminal microvilli of proximal progenitors and luminal budding of distal progenitors of the RV, along with the localization of tight junctions are in line with the luminal morphology and tight junctions we observed in proximal and distal tubules (Figures 4 and 6j). These findings show that ultrastructural analysis can provide additional insights into early cell type specification.

Later, in the SSB stage, cells did not possess microvilli, indicating their temporary disappearance in the process of further epithelialization. In the SSB, only podocytes had basal microprotrusions towards PECs (Figure 2c), a feature that has been proposed in adult mouse kidneys to serve as mechanosensing between podocytes and PECs or to signal changes in podocyte health.^31^ On top of these microprotrusions, podocytes developed microvilli starting in the CLS-1 (Figure 2), a feature in nephrogenesis that has not been described to our knowledge. It is conceivable that, in the process of urinary space formation and extensive changes in podocyte morphology, microvilli could have a mechanosensing role. Future studies may clarify the role of microvilli in other parts of the nephron and further the understanding of microvillus transformation in podocyte disease leading to nephrotic syndrome in adults.^32^ Apical microprotrusions and cilia were also noted in the cortical and medullary collecting duct (Figure 7c,g). This finding is consistent with earlier research of week 19 human kidney ultrastructure, where intercalated and principal cells possessed distinct apical protrusions.^10^ However, we could not determine with certainty any differences in number, location or size in protrusions between individual CD cells, indicating that intercalated and principal cells might not yet have developed at this stage. This interpretation is in line with transcriptomic data in which distinct CD cells (intercalated and principal cells) could only be discriminated after week 12 in human fetal kidneys.^33^

In addition to microvilli, we found that nuclear features can be informative to describe the maturity and cell type in the developing tubules. Previous studies have demonstrated that podocytes mature first (transcriptionally, in pseudo time) in the developing murine kidney followed by PT, LoH and DT.^34^ To our knowledge, it has not been shown which tubule segment matures first. However, ultrastructural features in our dataset indicate an order (Figure 6) and comparison to adult kidneys (Figure 4j) revealed the greatest similarity between proximal tubules of fetal and adult tissue, implying they mature first. Congruent features include the round, basal nuclei, a large cytoplasm creating space between individual nuclei, the brush border, and cytoplasmic lysosomes.

In contrast to these observations in PTs, morphological changes indicative of maturity over the course of the first trimester were less pronounced in DTs. DTs could be distinguished from collecting ducts by their location and absence of glycogen, which is present in large cytoplasmic aggregates in the CD (Figure 7). Fetal DTs are distinct from adult DTs in terms of their nuclear shape (raisin-shape instead of round), nuclear location (central instead of apical) and apical morphology (budding instead of smooth with few short micro protrusions) until week 12 (Figure 6i,j, Figure S6). These data indicate that the proximal tubule structurally matures before the distal tubule and that nuclear ultrastructural features can support the identification of immature tubules.

Nuclear morphology appears to have been an underappreciated feature in previous morphological studies of nephrogenesis despite many reports showing that nuclear shape is strongly linked to stem cell differentiation.^35^ Overall, stem cells, including their nuclei, are less rigid than mature cells and stiffen upon differentiation by an increase in the lamin A/C content leading to a decrease in chromatin remodeling.^35–39^ Consequently, nuclear shape and chromatin organization in stem cells and progenitor cells are distinct from differentiating and fully mature cells, and the nuclear shape is more likely to be affected by mechanical changes in the ECM.^36, 37^ In nephrogenesis, the formation of a lumen and thereby loss of apical cell–cell contact, sensation of luminal fluid flow, and the adhesion of perivascular cells on the basal side (Figure S7) could provide such mechanical cues. Although week 9 proximal tubular cells are not stem cells, it is likely that in their immature state they still possess a large amount of cellular plasticity. Consequently, it is possible that differences in nuclear shape are due to different cellular and nuclear stiffnesses and therefore distinct stages in differentiation.

In addition to nuclear shape, chromatin organization changed with the maturation of proximal tubular cells (Figure 4). A 3D reconstruction of stage 1 and 3 PT nuclei confirmed the morphological nuclear changes, from raisin-like to round shape (Figure 5). Furthermore, a proof of concept chromatin segmentation showed chromatin reorganization between stage 1 and stage 3 nuclei (Figure S3). Generally, it is believed that compacted heterochromatin is predominantly located in close proximity to the nuclear envelope and that it de-compacts in response to physiological stimuli and relocates into areas of the nucleus with high transcriptional activity.^40–42^ This relocalization and compaction is a cell type–dependent process. However, it appears to be consistent in a large variety of cell types including podocytes in mice.^43, 44^ To date, this has not been shown in other renal cell types and studies on human tissue are limited. Similarly, the nuclei of stage 3 compared to stage 1 proximal tubules in our dataset showed a lower amount of compacted chromatin in the nuclear periphery, potentially indicating the activation of a larger set of genes in the course of differentiation. However, additional techniques are needed to correlate changes in gene activity to morphological features. Nevertheless, the differences between stage 1 and stage 3 nuclei were large given the fact that the time between the stages was in the order of days. This is because the two stages could be detected in the same week 8 tissue and the developmental age difference between the tubules was correlated with anatomical location. Clearly, more drastic changes can be expected over a longer developmental time, particularly in terms of chromatin organization, as seen in neuron differentiation.^43^ Future studies could investigate chromatin organization and density combined with the location of cell type–specific genes that emerge early in nephrogenesis, to correlate morphological changes to maturity. New technological trends such as genome architecture mapping (GAM), a sequencing-based method that calculates statistical probability to determine the likelihood of physical interaction of two sequences in 3D based on cryosections,^45^ will facilitate such research.

To conclude, our analysis of the structural development of kidneys of the first trimester revealed important features that allow the determination of cell types at early stages in nephrogenesis and the order of maturation of proximal and distal segments. These findings are valuable to further understand nephrogenesis and to assess and improve engineered kidney tissue, such as kidney organoids, *in vitro*. Kidney organoids have been assessed structurally to a limited extent, likely due to the lack of human fetal kidney data as a reference and our research will support this in the future. Our study has also revealed new questions in need of further investigation to confirm the order of maturation of nephron segments and specific cell types, and to understand essential inter-communication and niches that need to be recreated in *in vitro* engineered tissue.

## Data sharing statement

All data generated for this article are made publicly available for download from https://doi.org/10.34894/SEHN8P, as well as readily accessible to view without downloading from the Image Data Resource (https://idr.openmicroscopy.org) under accession number idr0148.

## Acknowledgements

The authors wish to thank Dr Steven Lisgo and Berta Crespo Lopez from the Human Developmental Biology Resource (London) for their high-quality and quick work in providing the tissue for this study. The authors would also like to thank Dr Abdul Hamid from the pathology department of the MUMC for her initial opinion on the tissue health and Dr Virginie Joris for her technical support for Figure S2 and Figure S4. Finally, we would like to thank the Microscopy CORE Lab members of Maastricht University for their scientific and technical support.

