## Supplementary Figures for "Structural development of the human fetal kidney: new stages and cellular dynamics in nephrogenesis"

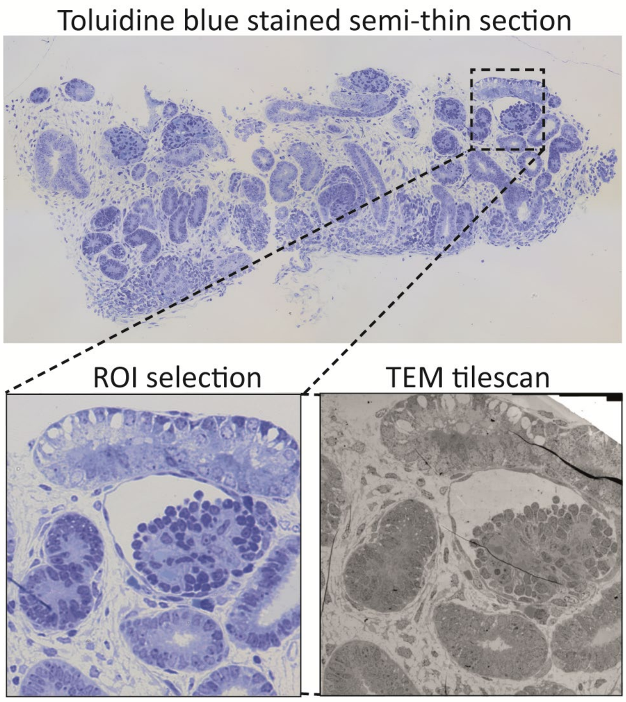


*Figure S1: Example of ROI selection in a toluidine blue stained semi-thin section and transmission electron microscopy (TEM) tile imaging of the selected ROI for ultrastructural assessment (patient ID 15333).*

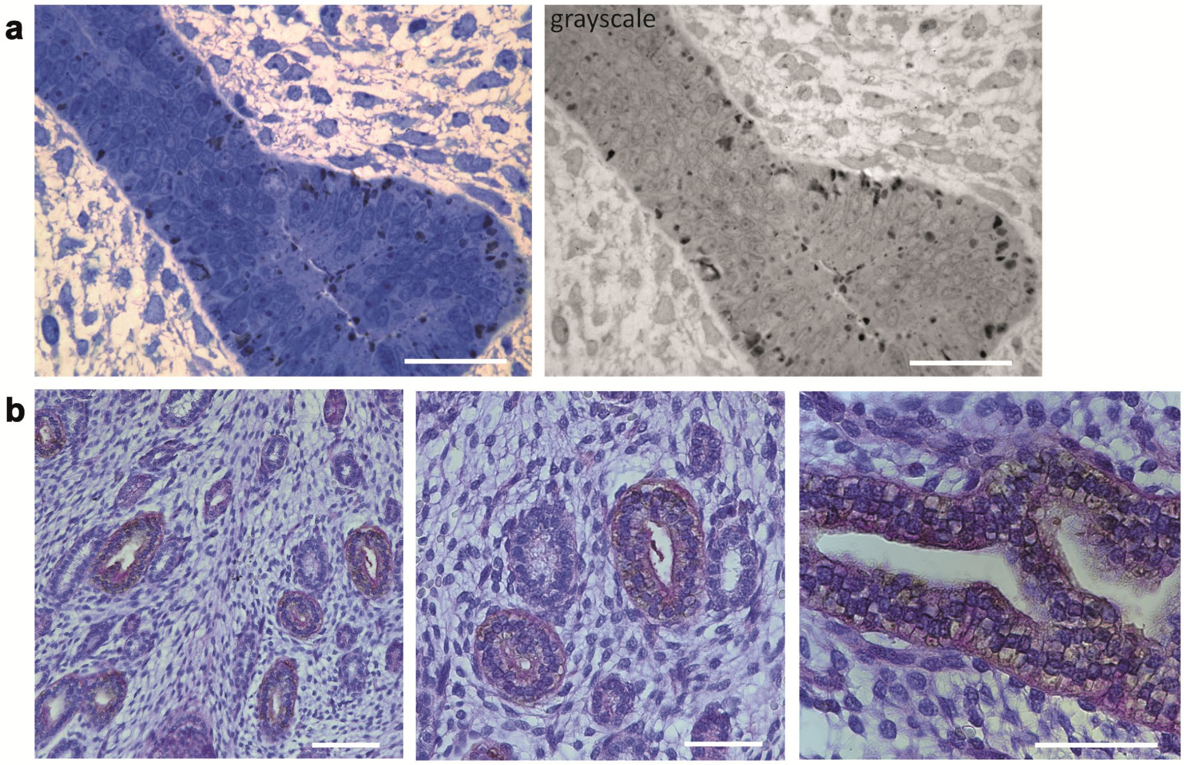


*Figure S2: a. Toluidine blue staining confirms dark-colored, dense glycogen aggregates in collecting duct cells (patient ID 15317). b. Periodic acid-Schiff staining (pink) and calbindin staining (brown) specific for ureteric buds. Scale bars: a: 30 µm, b: 50 µm.*

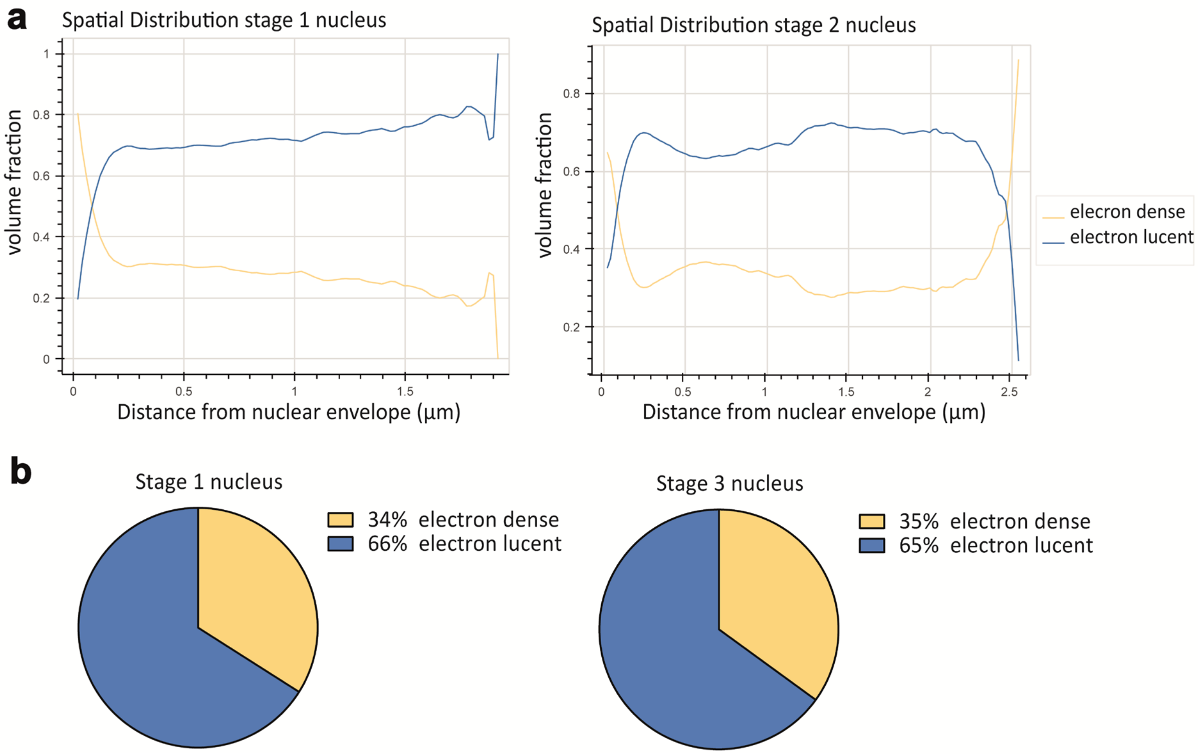


*Figure S3: Proof of concept quantification of chromatin in a stage 1 and a stage 3 proximal tubule. (a) Distance of the chromatin fraction from the nuclear envelope of electron-dense and electron-lucent chromatin. (b) Total fraction of electron-dense and electron-lucent chromatin per nucleus.*

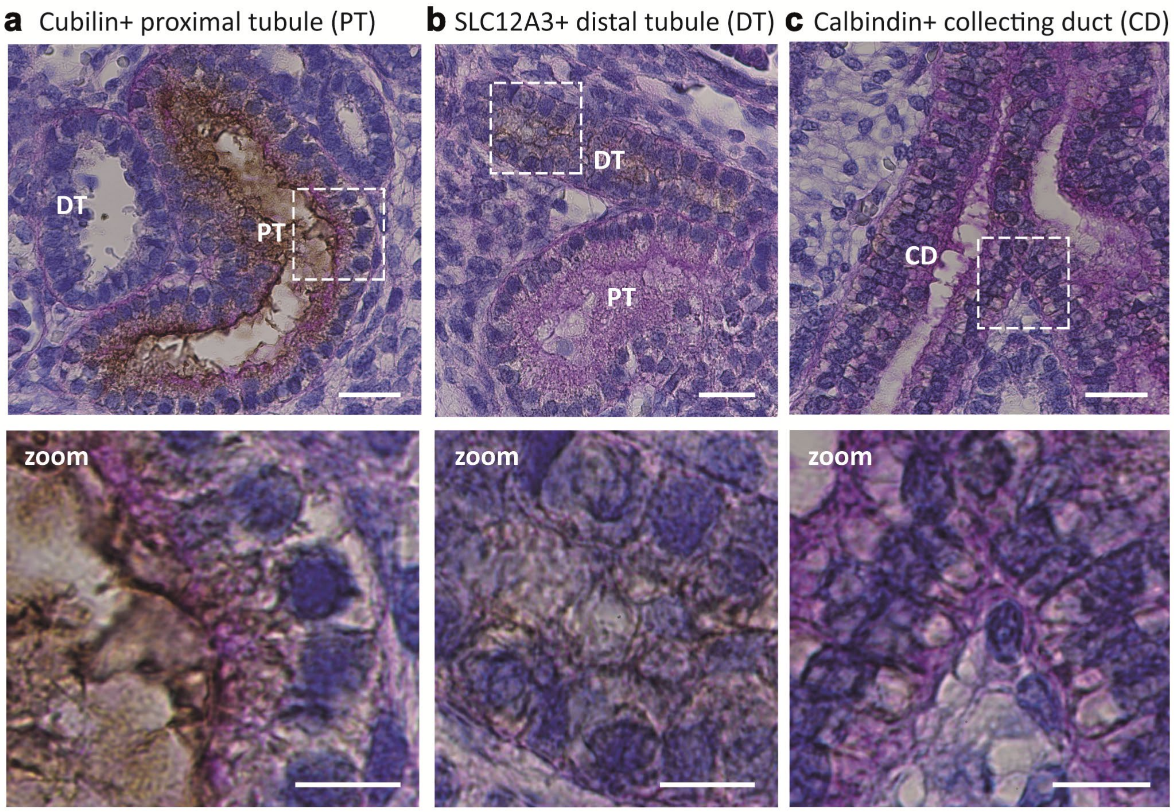


*Figure S4: Week 12 human fetal kidney (patient ID 15317) stained for (a) cubilin specific for proximal tubules (brown) (b) SLC12A3 specific for distal tubules (brown) and (c) calbindin specific for collecting ducts (brown). The sections are co-stained with Periodic acid-Schiff (pink) which predominantly stains the proximal tubules and collecting ducts. Scale bars upper row: 25 µm; scale bars zoom images: 10 µm.*

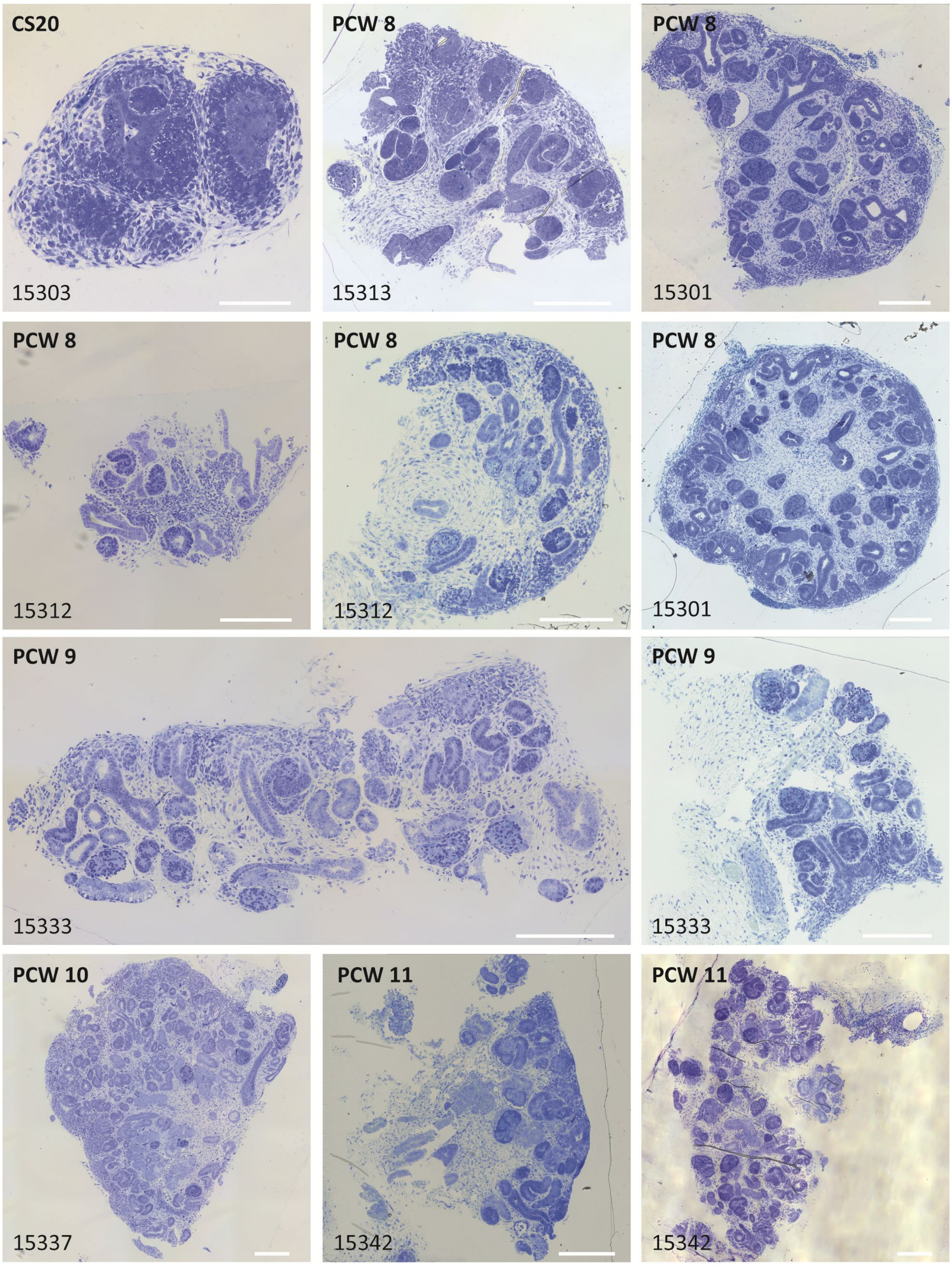
CS20
15303
PCW 8
15313
PCW 8
15301
PCW 8
15312
PCW 8
15312
PCW 8
15301
PCW 9
15333
PCW 9
15333
PCW 10
15337
PCW 11
15342
PCW 11
15342

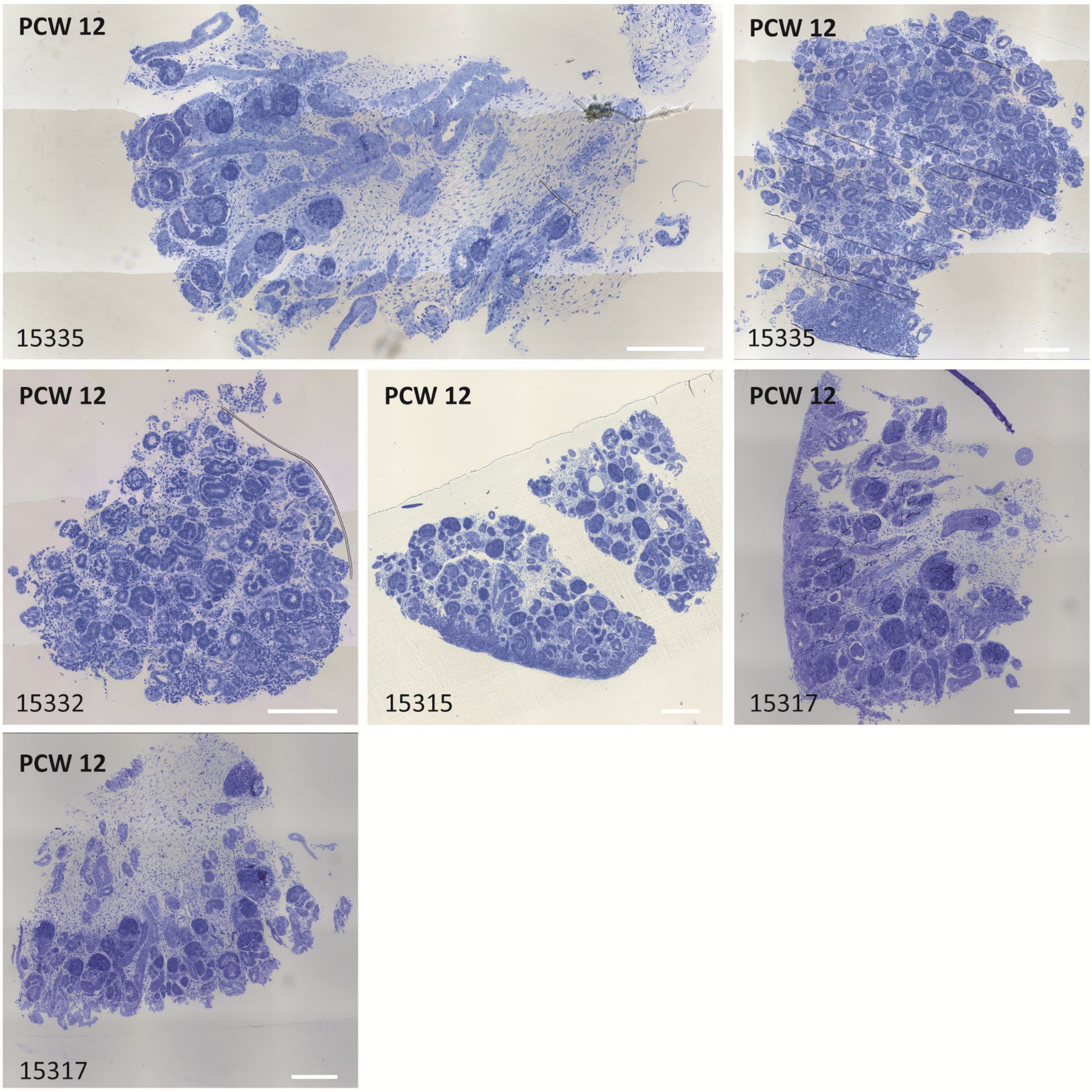


*Figure S5: Toluidine blue stained semi-thin sections of all fetal kidneys included in this study and from which ROIs for large TEM tile scans were chosen. Scale bars: CS20: 100 µm, all other scale bars: 200 µm.*

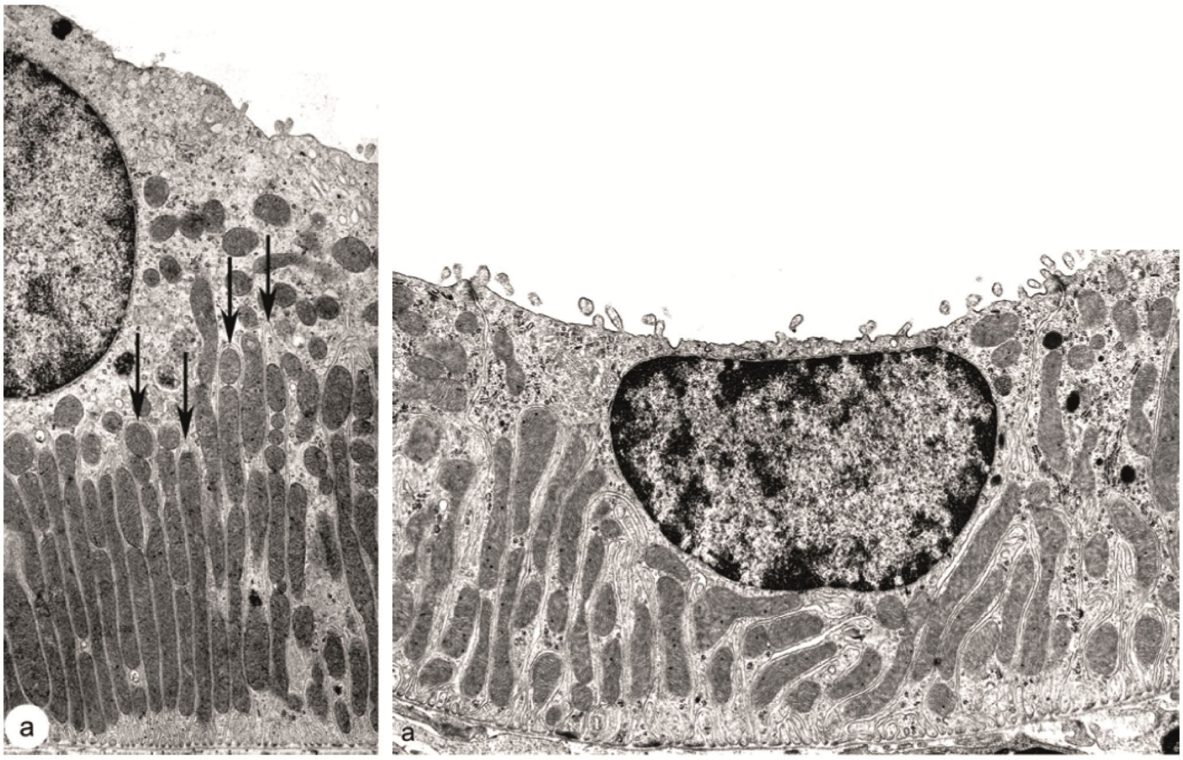


*Figure S6: Cross-sections of adult rat distal tubules, adapted from Kriz and Kaissling [20].*

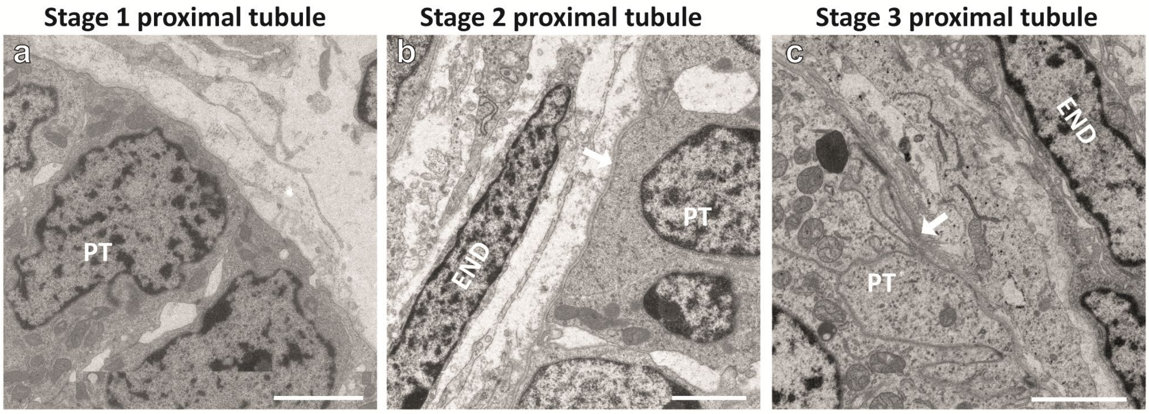


*Figure S7: Stages of proximal tubule (PT) development described in Figure 4 show an increase in the vascular attachment to the PT. (a) The peritubular vasculature remains distanced from the PT in the interstitium. (b) The peritubular vasculature lined with endothelial cells (END) in close proximity to PT. (c) The peritubular vasculature adheres to the basement membrane (white arrow) of the PT. Scale bars: a, c: 3 µm, b: 2 µm.*
