## Supplementary Methods for "Structural development of the human fetal kidney: new stages and cellular dynamics in nephrogenesis"

### Tissue embedding and processing for TEM

Tissue that was assessed as healthy by PAS-stained sections was used for further TEM analysis. The kidneys were cut into 1 $mm^3$ cubes using sharp razor blades while immersed in Karnovsky's fixative (2.5% glutaraldehyde (Merck) and 2% paraformaldehyde (PFA, Sigma-Aldrich) in 0.1 M sodium cacodylate (Acros), pH 7.4) overnight at 4°C. After washing with 0.1 M cacodylate buffer, the samples were postfixed with 1% osmium tetroxide (Agar Scientific) in the same buffer containing 1.5% potassium ferricyanide (Merck) for 1 h in the dark at 4°C. Next, the samples were rinsed with Milli-Q water, and dehydrated at room temperature (RT) in a graded ethanol series (70, 90, 100%, VWR), infiltrated with Epon resin (LADD), embedded in the same resin, and polymerized for 48 h at 60°C. The region of interest (ROI) was determined in semi-thin sections (1 µm thick) stained with toluidine blue (Merck) (Figure S1). Ultrathin sections of 60 nm were subsequently cut using a diamond knife (Diatome) on a Leica UC7 ultramicrotome, and transferred onto 50 mesh copper grids covered with a Formvar and carbon film. The sections were then stained with 2% uranyl acetate (SPI) in 50% ethanol (VWR) and Reynold's lead citrate (Merck).

### 3D nuclear reconstruction and chromatin segmentation

Individual images were each registered with the previous image and transformed to create the aligned image stack using pystackreg.[17] The aligned stack was 3D-median filtered before segmentation in Amira 6.5 (Thermo Fisher Scientific).[18] The nucleus was manually segmented and border pixels of the nuclear envelope were removed. Two classes of chromatin were identified: the electron-dense chromatin class and the electron-lucent class. They were segmented via thresholding after masked histogram equalization. To visualize the spatial distribution of the chromatin classes, the shortest Euclidian distances between chromatin pixels and nuclear envelope pixels were measured using scikit-image.[18] For stage 1 and stage 3 proximal tubules, one representative nucleus was segmented for 3D visualization of TEM findings as a proof of concept.

Combined immunostaining and Periodic Acid-Schiff (PAS) staining

Paraffin sections of week 12 human fetal kidneys were deparaffinized using Neoclear (EMD Millipore) and rehydrated in an ethanol series. The sections were then incubated in citrate buffer (0.01 mol/L) in a waterbath (1 hour, 95°C), after which they were cooled down during 15 minutes to RT. Inhibition of endogenous peroxidases was done by incubating the sections with 3% $H_2O_2$ for 20 minutes at RT. After blocking with 5% goat serum and 5% bovine serum albumin in TBS-Triton 0.05% for 1 hour at RT, the sections were incubated overnight at 4°C with the appropriate antibody: monoclonal anti-calbindin antibody (C9849, Sigma-Aldrich) diluted 1/40, monoclonal anti-cubilin antibody (ab191073, Abcam) diluted 1/300, polyclonal anti-SLC12A3 antibody (HPA028748, Sigma-Aldrich) diluted 1/200. After washing three times with PBS-Triton 0.05%, the sections were incubated for 1h at RT with the respective goat anti-mouse (1706516, Bio-Rad) or goat anti-rabbit (1706515, Bio-Rad) – HRP conjugated antibody diluted 1/300. After additional washing, the sections were treated with a diaminobenzidine (DAB) substrate kit (Abcam) according to the manufacturer’s protocol. Counterstaining was performed using a Periodic acid Schiff kit (Sigma-Aldrich) according to the manufacturer’s protocol. After dehydration in an ethanol series and additional incubation in Neoclear, the slides were mounted with Ultrakitt (VWR). The slides were imaged using an automated Nikon Eclipse Ti-E microscope equipped with a 60× oil objective.
